# Screening Avian Pathogens in Eggs from Commercial Hatcheries in Nepal- an Effective Poultry Disease Surveillance Tool

**DOI:** 10.1101/2022.08.11.503567

**Authors:** Shreeya Sharma, Kavya Dhital, Dhiraj Puri, Saman Pradhan, Udaya Rajbhandari, Amit Basnet, Sajani Ghaju, Prajwol Manandhar, Nabin U Ghimire, Manoj K Shahi, Ajit Poudel, Rajindra Napit, Dibesh Karmacharya

**Author notes:** these authors contributed equally (Shreeya Sharma, Kavya Dhital).

## Abstract

**Background:** Commercial hatcheries play an important role in the overall poultry value chain-providing small to large poultry farmers with day old chicks. Any outbreak in such hatcheries can spread diseases to other farms. Regular screening of major avian pathogens, along with strict bio-security measures, can prevent spread of diseases in hatcheries. Newcastle Disease Virus (NDV), Infectious Bronchitis Virus (IBV), *Mycoplasma gallisepticum* (MG), *Mycoplasma synoviae* (MS), Infectious Bursal Disease Virus (IBDV) and Influenza A Virus (IAV) are among the most prevalent poultry diseases which can be detected in egg albumin.

**Method:** We retrospectively (August 2020-August 2021, except October 2020) analyzed diagnostic results for six selected avian pathogens (NDV, IBV, MS, MG, IBDV and IAV) on eggs (n=4343) received from eleven major commercial poultry hatcheries located in the five adjoining districts of Kathmandu, Nepal. Albumin from 10% randomly selected eggs from each hatchery were tested for the six avian pathogens using multiplex PCR.

**Result:** Majority (7/11, 64%) of the poultry hatcheries had at least one of the six pathogens present. We detected at least one avian pathogen in nine out of eleven months (82%) of screening. Except for IBDV, we found one or more of the other major avian pathogens-Influenza A (IAV) (n=4 times) and *Mycoplasma gallisepticum* (MG) (n=4 times) were detected the most, followed by Newcastle Virus (NDV) (n=3 times). Infectious bronchitis virus (IBV) were detected twice, and *Mycoplasma synoviae* (MS) was detected once.

**Conclusion:** In a resource strapped country like Nepal, poultry disease outbreak investigation in particular and surveillance in general are challenging. Meanwhile, poultry production is highly impacted by disease outbreaks often triggered by poor bio-security and lack of pathogen screening practices. Our molecular screening tests have picked up major poultry pathogens present throughout the year in eggs collected from hatcheries. Influenza A was detected at 4 different incidences throughout the year, which is of concern to both human and animal health. Quick systematic screening of eggs at key distribution points (hatcheries) for major avian pathogens is an effective surveillance tool for early disease detection and containment of outbreaks.

## Introduction

Globally, the poultry sector is a sizeable industry with a current market value of $ 310.7 billion and is expected to grow at a compound annual growth rate (CAGR) of 3.8%^1^. Poultry is a rapidly growing agricultural sub-sector in developing countries^2^, however, product quality, safety, and avian diseases continue to be a major challenge to this industry^3^.

Hatcheries occupy a focal position in the poultry production chain, connecting with multiple flocks^4^, thereby acting as a reservoir, linkage and source of pathogenic microorganisms^5^. Nepal’s $240-million poultry industry^6^ is buttressed by 21,956 poultry farms present in sixty four out of seventy five districts, where 325 total commercial hatcheries represent this burgeoning industry^7^.

Animal trade related movement of poultry and poultry products from production sites, such as hatcheries, can influence disease transmission dynamics into uncontaminated flocks^8^. For example, transmission of a recent subtype of Avian Influenza virus in Bangladesh was associated with poultry movement^9^.

Several pathogens (both mono and multi-causal) have been implicated as probable causes of avian diseases. Poultry can be infected or colonized with other potential organism via eggs^3^. Contaminated eggs can be a source of infection and a vehicle for transmission of pathogens ^10, 11^.Contamination can occur horizontally through egg shells^12^ or vertically before oviposition stemming from infection of reproductive organs^13^. In vertical/ trans-ovarian route, the disease is ascendingly transmitted from laying hen to its progenies-where the yolk, albumen and membranes are contaminated via the reproductive organs^14^ before the eggs are covered by shell in the uterus. Handling fecal material, dust, and dirt can contaminate eggs in hatcheries through horizontal route. Extrinsic factors such as temperature, moisture, shell characteristics and membrane properties are attributable to pathogen transmission^15^.

Eggs form as great a proportion of the animal protein diet for Low and Middle Income Countries (LMICs), overlooking a projected 76.6%^16^ growth in egg production. Egg production is imperative to the growing population for providing an inexpensive source of protein^17^, thereby contributing to food security. Global egg production continues to see substantial growth from 61.7 tons to 76.7 tons^18^, a 24% increase in the past decade. Asia is the largest egg producing region, contributing to 60% of the total production volume^16^.

Avian pathogens can cause huge economic loss (>20%) in the overall poultry production, and three times due to loss from mortality^19, 20^. Egg and egg based product surveillance programs are highly effective in controlling foodborne disease outbreaks-often providing information for timely intervention, control and mitigation measures^21, 22, 23^. Egg-based surveillance helped identify more than 895 foodborne disease outbreaks in Spain (2000-2002), majority (85%) caused by *Salmonella* ^24^.

Most studies have focused on detecting of foodborne pathogens like *Salmonella* spp., *Camphylobacter* spp. and *Escherichia coli* in eggshells^26,27,28^, we posit that albumin-based screening is also a convenient tool and useful in detecting other important avian pathogens such as-*Mycoplasma gallisepticum* (MG), *Mycoplasma synoviae* (MS), Infectious Bronchitis Virus (IBV), Influenza A Virus (IAV), Newcastle Disease Virus (NDV) and Infectious Bursal Disease Virus (IBDV) in hatcheries to minimize contamination through horizontal and vertical transmission modes. Avian pathogens have been isolated from oral swabs, cloacal swabs, serum samples, egg yolk, egg shells, and environmental swabs but albumin-based molecular detection has not been intensively used till date. Due to the dearth of literature available on albumin screening, we used Polymerase Chain Reaction (PCR) based tests to screen for six major avian pathogens (IBD, IBDV, MS, MG, IAV and NDV) in egg albumin from eggs collected from eleven hatcheries, hence devising cost-effective poultry pathogen surveillance tool in hatcheries.

## Selected Avian Diseases

### *Mycoplasma synoviae* and *Mycoplasma gallisepticum*

Mycoplasma is a vertically transmitted disease^29^ with pronounced effects in eggshell-altered surface, thinning, translucency, consequently leading to a greater incidence of eggshell cracks and breaks^30^. Though it is a non-fatal disease^31^, it can significantly affect weight gain, feed conversion ratio, fertility, chronic respiratory disease and hatchability^32, 33^ in birds. MG and MS are bacterial OIE-listed respiratory pathogens^34^ which often persist in sub-clinical level^35^ and are a key cause for economic loss in the poultry industry^34^. Mycoplasma infections, especially in farms with weak biosecurity, are often the cause of eggshell abnormalities and decrease in egg production^36^.

### Newcastle Disease Virus (NDV)

Newcastle disease (ND), an OIE-notifiable List A disease, is caused by avian paramyxovirus serotype 1 (APMV-1) virus^38^ of Avulavirus genus. It is one of the highly pathogenic viral diseases of avian species, and a major cause of morbidity and mortality in flocks^39^. Affected birds develop respiratory, digestive and neurologic symptoms with profound immunosuppression^40^. In many countries throughout Asia and Africa, ND remains endemic in commercial poultry despite intensive vaccination program that have been applied for decades^41^. NDV can replicate in the reproductive tract of hens and contaminate internal components of eggs and eggshell surface^42^.

### Infectious Bronchitis Virus (IBV)

Infectious bronchitis in poultry is caused by IBV-an Avian Coronavirus (ACoV) of genus Gammacoronavirus^43^. IBV causes a fast-spreading respiratory disease in young chicks, with laying hens experiencing reduced production, egg shell abnormalities, and decreased internal egg quality^44^. Along with commercial poultry, backyard poultry and free-ranging birds may serve as ‘reservoir’ for ACoV transmission, and migratory birds often acting as an intermediary host spreading to wide and distant areas^45^.

### Infectious Bursal Disease Virus (IBDV)

Infectious Bursal Disease, commonly known as Gumboro, is an immunosuppressive disease transmitted mainly horizontally through the feco-oral route^46^. It is caused by a double stranded RNA virus-IBDV (genus *Avibirnavirus* of family *Birnaviridae*)^47^. There are two distinct serotypes of the virus, but only serotype 1 viruses cause disease in poultry^48^. Viruses belonging to one of these antigenic subtypes are commonly known as variants, causing up to 60 to 100 percent mortality rates in chickens^49^.

### Influenza A virus (IAV)

Avian Influenza (AI), caused by IAV, is a highly contagious viral infection which may cause up to 100% mortality in domestic chickens or turkeys^50^. The disease is caused by a highly mutable RNA virus that belongs to the family *Orthomyxoviridae*^51^. Influenza viruses have two surface proteins, hemagglutinin (HA) and neuraminidase (NA)^52^ that determine their subtype and the animal species they infect; there are 16 HA and nine NA types^53^. When AI viruses of two HA types, H5 and H7, infect domestic poultry (chickens and turkeys) they often mutate and virulent disease arises in these birds which is called highly pathogenic avian influenza (HPAI)^54^. The initial infection that causes subclinical or mild disease is called low pathogenic avian influenza (LPAI)^55^. Wild water birds act as reservoir hosts of IAV, however these viruses generally do not cause disease in these birds^56^.

## Methodology

We retrospectively looked at the data on the presence of 6 major avian pathogens on eggs received periodically every month (August 20-August 2021, except October 2020) from the eleven major hatcheries in and around Kathmandu valley. These hatcheries participated in preventive pathogen screening services provided by the BIOVAC Nepal’s Poultry Diagnostic Laboratory (BNPDL). The sampling was performed by trained personnel. No live animals were harmed and the study does not include handling of animal. No embryonated eggs were killed during sampling process-qualifying this study to be exempted from any ethical approval. To maintain anonymity of the hatcheries, they were coded with numeric digits on the basis of time the samples were received. A total of 4343 eggs from eleven major hatcheries located in the five surrounding districts (Kathmandu, Bhaktapur, Lalitpur, Kavrepalanchowk and Ramechhap) of Kathmandu, Nepal were received and tested (Figure 1).

**Figure 1:**
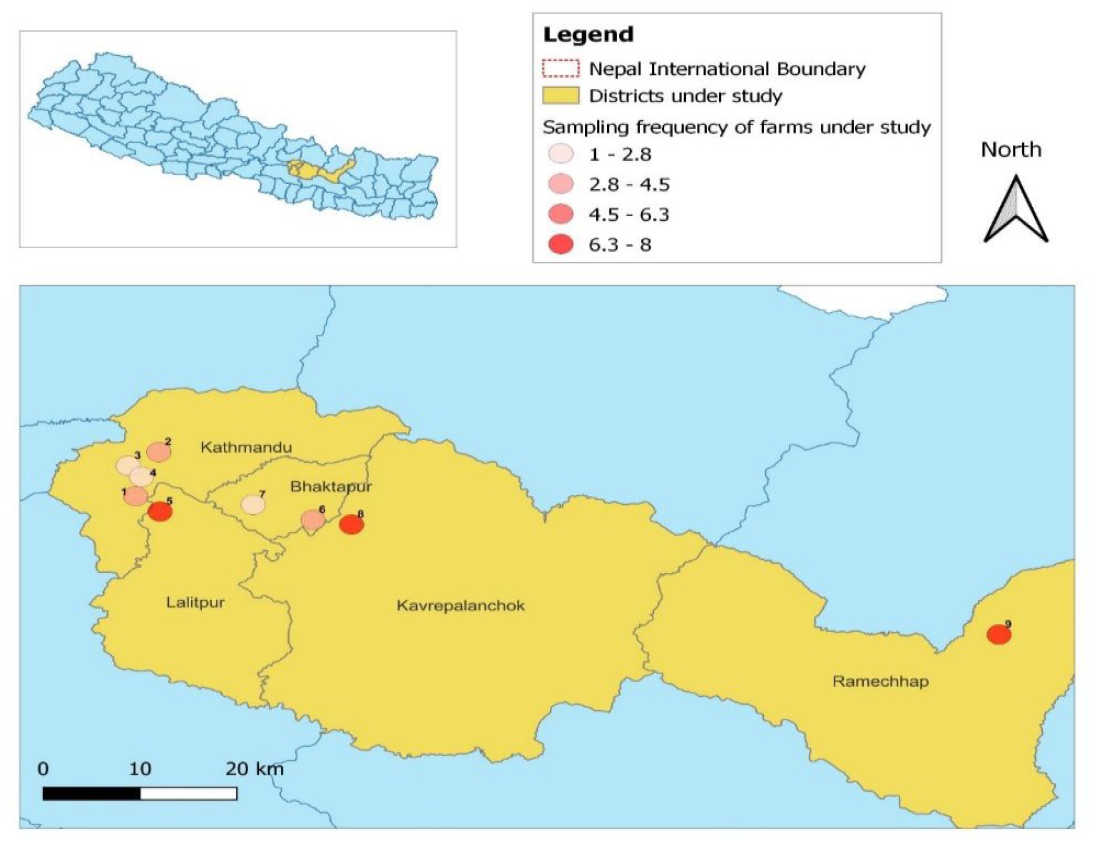
Participating eleven major hatcheries located in Kathmandu and surrounding five districts. (Kathmandu, Bhaktapur, Lalitpur, Kavrepalanchowk and Ramechhap). As part of preventive disease screening, eggs are routinely received by BIOVAC Nepal’s Poultry diagnostic laboratory located in Banepa (Nala), Nepal.

These eggs were brought to the BNPDL every month (except October 2020) in batches (133±60 eggs per batch) packaged in crates (30 eggs per crate). Albumin extracted from 10% random eggs from each batch (3 eggs from each crate) were tested for six selected Avian pathogens (NDV, IAV, IBV, IBDV, MS and MG) using PCR (Figure 2)

**Figure 2:**
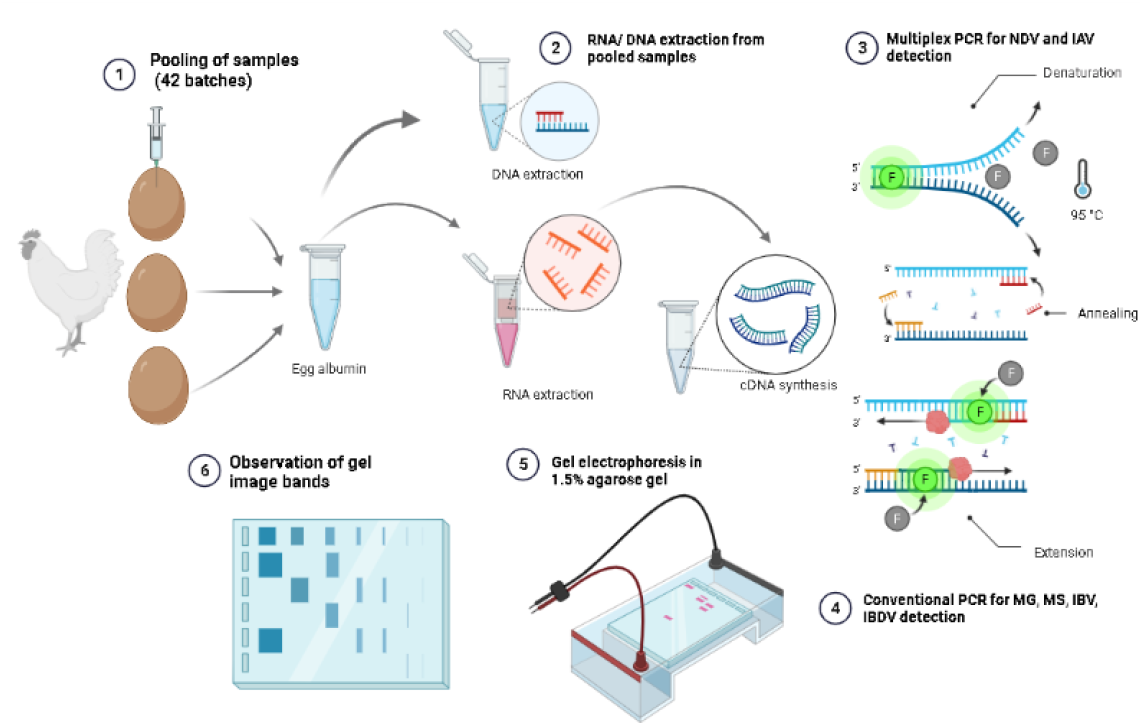
Laboratory testing for the detection of six avian pathogens: BIOVAC Nepal’s Poultry Diagnostic Laboratory (BNPDL) received samples from eleven participating hatcheries for preventive diagnostic screening of avian pathogens.

### Molecular Detection of Six Avian Pathogens

#### Nucleic Acid Extraction and cDNA synthesis

The nucleic acids (DNA/RNA) from pooled egg albumin samples were extracted using automated nucleic acid extractor (abGenix™ AITbiotech, Singapore) following manufacturer’s instructions. cDNA for the extracted nucleic acids were synthesized using iScript™ cDNA Synthesis Kit (Bio-Rad Laboratories, USA). For a single sample, 4 μL of 5X iScript reaction mix, 1 μL of iScript reverse transcriptase, 6 μL of nuclease free water and 9 μL of the extracted nucleic acid was used to prepare 20 μL of cDNA. The cDNA was synthesized in thermal cycler by incubating the mix at 25°C for 5 minutes followed by reverse transcription at 46°C for 20 minutes and RT inactivation at 95°C for 1 minute. PCR for IBV, IBD, MG and MS were performed using QIAGEN Multiplex PCR Kit (Qiagen, Catalog No. 206145). Multiplex PCR was used to detect IAV and NDV simultaneously in the samples using QIAGEN Multiplex PCR kit.

#### Multiplex PCR for IAV and NDV

We have developed and optimized a multiplex PCR that detects both IAV and NDV simultaneously in one single test. A 291 bp fragment of Matrix protein gene of NDV and 156 bp fragment of Matrix protein gene of IAV was amplified in 25 μL of the reaction mixture containing: 3 μL of cDNA, 5 μL QIAGEN® nuclease-free water, 2.5 μL of 5X Q Solution, 0.5 μL NDV primer (forward), 0.5 μL NDV primer (reverse), 0.5 μL IAV primer (forward), 0.5 μL IAV primer (reverse) and 12.5 μL of 2X of QIAGEN® Multiplex PCR Master Mix (HotStarTaq DNA Polymerase, MgCl2, dNTPs and PCR buffer). PCR condition: 1 cycle of initial denaturation at 95°C for 15 minutes, 45 cycles of denaturation at 95°C for 20 seconds, annealing at 60°C for 20 second and extension at 72°C for 30 second. The PCR ended with a final elongation at 72°C for 5 minutes. PCR products were visualized in Gel electrophoresis (1.5%) (Figure 3).

**Figure 3:**
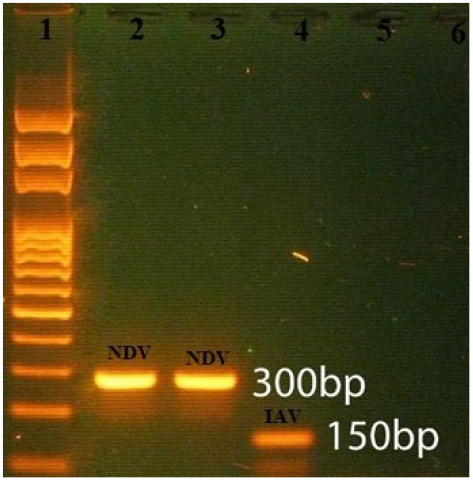
Detection of Influenza A (IAV) and Newcastle Disease Virus (NDV) using Multiplex PCR: Newcastle Disease Virus (NDV) is detected as 300bp PCR amplicon, and Influenza A Virus (IAV) as 150bp. Lane 1: DNA ladder; Lanes 2-5: pooled albumin samples, Lane 6: negative control. Visualized under 1.5% gel electrophoresis.

#### PCR Primers-multiplex IAV and NDV

For NDV, 10 pico-molar concentration each of forward primer (5’-GCTCAATGTCACTATTGATGTGG-3’) and reverse primer (5’-TAGCAGGCTGTCCCACTGC-3’) were used and for IAV, 10 pico-molar concentration each of forward (5’-CTTCTAACCGAGGTCGAAACG-3’) and reverse (5’GGTGACAGGATTGGTCTTGTC-3’) were designed using NCBI PrimerBlast®.

#### PCR detection of IBV

A 433 bp fragment of 3’ UTR of IBV was amplified in 25 μL of the reaction mixture containing: 2 μL of template cDNA, 8.5 μL QIAGEN® nuclease-free water, 1 μL All 1-F primer (forward), 1 μL Del1-R primer (reverse) and 12.5 μL of 2X of QIAGEN® PCR Master Mix. PCR condition: 1 cycle of initial denaturation at 95°C for 15 minutes, 35 cycles of denaturation at 95°C for 30 seconds, annealing at 57°C for 30 second and extension at 72°C for 40 second. The PCR ended with a final elongation at 72°C for 5 minutes.

#### PCR Primers-IBV

10 pico-molar concentration of each All 1-F forward primer (5’-CAGCGCCAAAACAACAGCG-3) and Del1-R reverse primer (5’-CATTTCCCTGGCGATAGAC-3’) were used for detection of IBV as per Saba et al. (2014)^57^.

#### PCR detection of IBDV

A 643 bp fragment of complete hyper variable region of VP2 gene of IBDV was amplified in 25 μL of the reaction mixture containing 2 μL of template cDNA, 8.5 μL QIAGEN® nuclease-free water, 1 μL Infectious Bursal Disease Forward Primer, 1μL Infectious Bursal Disease Forward Primer and 12.5 μL of 2X of QIAGEN® PCR Master Mix. PCR conditions: 1 cycle of initial denaturation at 95°C for 15 minutes, 35 cycles of denaturation at 95°C for 30 seconds, annealing at 53°C for 20 second and extension at 72°C for 45 second. The PCR ended with a final elongation at 72°C for 5 minutes.

#### PCR Primers-IBDV

10 pico-molar concentration of each forward primer (5’-TCACCGTCCTCAGCTTAC-3’) and reverse primer (5’-TCAGGATTTGGGATCAGC-3’) were used for the detection of IBD as per Kataria et al. (2007)^58^.

#### PCR detection of MS

A 207 bp fragment of 16s rRNA gene of MS was amplified in 25 μL of the reaction mixture containing 2 μL of template cDNA, 8.5 μL QIAGEN® nuclease-free water, 1 μL Mycoplasma synoviae forward primer, 1 μL Mycoplasma synoviae reverse primer, and 12.5 μL of 2X of QIAGEN® PCR Master Mix. PCR conditions: 1 cycle of initial denaturation at 95°C for 15 minutes, 35 cycles of denaturation at 95°C for 30 seconds, annealing at 53°C for 20 second and extension at 72°C for 15 second. The PCR ended with a final elongation at 72°C for 5 minutes.

#### PCR Primers-MS

10 pico-molar concentration of each forward primer (5’-GAGAAGCAAAATAGTGATATC-3’) and reverse primer (5’-TCGTCTCCGAAGTTAACAA-3’) were used for detection of MS as per Kahya et al. (2015)^59^.

#### PCR detection of MG

A 185 bp fragment of 16s rRNA gene of MG was amplified in 25 μL of the reaction mixture containing 2 μL of template cDNA, 8.5 μL QIAGEN® nuclease-free water, 1 μL *Mycoplasma gallisepticum* forward primer, 1 μL *Mycoplasma gallisepticum* reverse primer and 12.5 μL of 2X of QIAGEN® PCR Master Mix. PCR conditions: 1 cycle of initial denaturation at 95°C for 15 minutes, 35 cycles of denaturation at 95°C for 30 seconds, annealing at 53°C for 20 second and extension at 72°C for 15 second. The PCR ended with a final elongation at 72°C for 5 minutes.

#### PCR Primers-MG

10 pico-molar concentration each of forward primer (5’-GAGCTAATCTGTAAAGTTGGTC-3’) and reverse primer (5’-GCTTCCTTGCGGTTAGCAAC-3’) were used for detection of MG as per Kahya et. al. (2015).

All PCR amplified products were visualized under 1.5% agarose gel electrophoresis.

## Results

We retrospectively looked at the data on the presence of 6 major avian pathogens on eggs received periodically every month (August 20-August 2021, except October 2020) from the eleven major hatcheries in and around Kathmandu valley. These hatcheries participated in preventive pathogen screening services provided by BNPDL. The hatcheries had experienced high morbidity and mortality in their birds; and had seen decreased and defective egg production.

In an average we received 430 eggs every month from one or more of the eleven hatcheries, majority (7/11, 64%) had at least one of the six pathogens present. We detected at least one avian pathogen in nine out of eleven months (82%) of screening. Except for IBDV, we found one or multiple occurrence of other major avian pathogens-Influenza A (IAV) (n=4 times) and *Mycoplasma gallisepticum (MG)* (n=4 times) were detected the most, followed by Newcastle Virus (NDV) (n=3 times). Infectious bronchitis virus (IBV) were detected twice, and *Mycoplasma synoviae* (MS) was detected once (Table 1).

**Table 1:** Eggs received and avian pathogen detected from each hatchery: Between Aug 2020 through August 2021 (except October 2020), we received eggs from participating 11 major poultry hatcheries. Eggs received from some hatcheries (1, 2, 3 and 7) were free of screened avian pathogens. Influenza A (IAV) (n=4 times) and Mycoplasma gallisepticum (MG) (n=4 times) were detected the most, followed by Newcastle Virus (NDV) (n=3 times). Infectious bronchitis virus (IBV) were detected twice, and Mycoplasma synoviae (MS) was detected once.

| HATCHERY | Months |  |  |  |  |  |  |  |  |  |  |  |
| --- | --- | --- | --- | --- | --- | --- | --- | --- | --- | --- | --- | --- |
|  | Aug 20 | Sep 20 | Nov 20 | Dec 20 | Jan 21 | Feb 21 | Mar 21 | Apr 21 | May 21 | Jun 21 | Jul 21 | Aug 21 |
| 1 | N=260 |  |  |  |  |  |  |  |  |  |  |  |
| 2 | N=120 |  |  |  |  |  |  |  |  |  |  |  |
| 3 |  | N= 450 |  |  |  |  |  |  |  |  |  |  |
| 4 |  | N=150<br>IAV |  | N=120<br>IAV/IBV |  |  |  |  |  |  |  |  |
| 5 |  |  | N= 60 | N=240<br>NDV |  |  |  |  |  |  |  |  |
| 6 |  |  |  |  | N=420<br>NDV |  |  |  |  |  |  |  |
| 7 |  |  |  |  |  | N=60 | N=210 |  |  |  |  |  |
| 8 |  |  |  |  |  | N=27<br>MS IBV |  |  |  |  |  |  |
| 9 |  |  |  |  |  |  | N=65 | N=30<br>MG | N=180<br>MG | N=300<br>MG, NDV, IAV | N=418 |  |
| 10 |  |  |  |  |  |  |  |  | N=10 | N=270 | N=870<br>MG |  |
| 11 |  |  |  |  |  |  |  | . |  |  | N=90 | N=1050<br>IAV |

In hatchery 4, we detected IAV in samples received in two separate months (September and December 2020). Meanwhile, we received most consecutive samples from hatchery 9, where we detected MG 3 months in a row (April, May and June 2021), with multiple pathogens (MG, IAV and NDV) present in June 2021.

In the winter season (January-April 2021), four batches had four detectable pathogens (NDV, IBV, MS and MG). In rainy or wet season (May-August 2021), three different pathogens (IAV, MG and NDV) were found in the 9 batches; during this period MG was detected in three consecutive batches. During the fall season (September-December), we detected only three pathogen (NDV, IBV and IAV) in four batches. We detected more pathogens during rainy or wet season than in winter or fall season (Figure 4).

**Figure 4:**
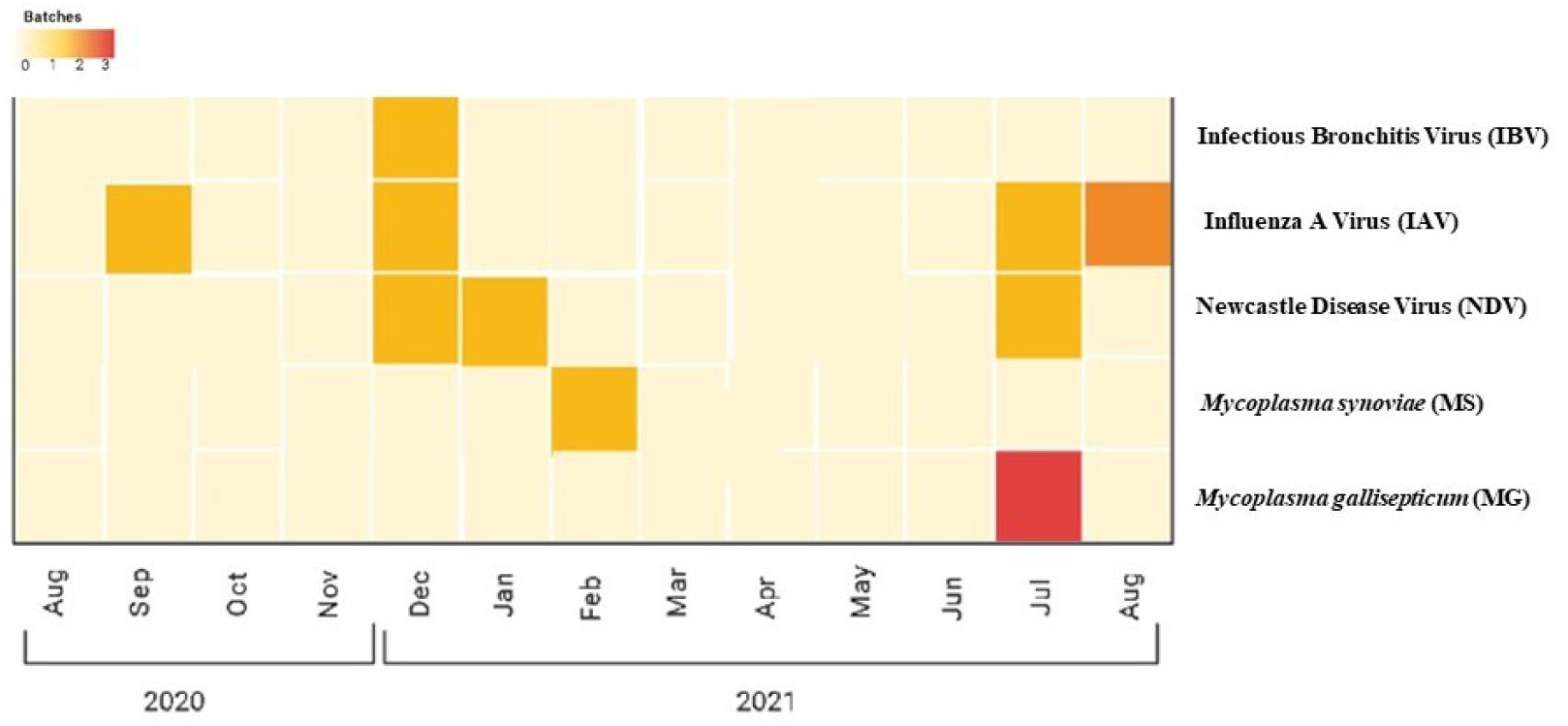
Detected avian pathogens in received batches from the eleven different hatcheries: In the winter season (January-April 2021), four batches had four detectable pathogens (NDV, IBV, MS and MG). In rainy or wet season (May- August 2021), three different pathogens (IAV, MG and NDV) were found in the 9 batches; during this period MG was detected in three consecutive batches. During the fall season (September-December), we detected only three pathogen (NDV, IBV and IAV) in four batches. We detected more pathogens during rainy or wet season than in winter or fall season (Figure 3).

## Discussion

There is no active surveillance of most of the avian pathogens in Nepal, only outbreak related Avian Influenza (Bird flu) is investigated^60, 61^ primarily due to human health concerns. Meanwhile, disease outbreaks in poultry farms are often reported based on clinical symptoms and mortality. Lack of comprehensive animal diagnostic facilities also limits the farmers’ ability to have disease outbreaks properly investigated. Preventive disease screening is still a novel concept in Nepal’s poultry industry. Against that backdrop, we have set up a poultry disease diagnostic laboratory (BNPDL) based in Kathmandu and Nala, Nepal. We routinely receive requests to screen for avian pathogens from poultry farmers, including hatcheries. Hatcheries can be a source of disease spread through contaminated eggs and day old chicks. We have detected presence of avian pathogens in egg albumin using molecular (PCR) detection. In a resource strapped country like Nepal, where disease surveillance is not well developed, a routine egg screening based on accurate and relatively fast PCR screening, can offer an important insight into floating avian pathogens in poultry population at any given time. This kind of information can be helpful for poultry producers, including hatcheries, to prevent and mitigate their losses by adopting appropriate interventions.

### Egg-shell based pathogen detection

There are some egg-shell based poultry disease surveillance program being used in some countries, however, they have only focused on food borne pathogens such as *Salmonella, Camphylobacter* and *Escherichia coli*^62,63,64,65^. Majority (41%) of all foodborne disease outbreaks in Spain were associated with consumption of eggs and egg products^66^, which explains a high level of contamination associated with eggs. Similarly, quarterly surveillance measures have also been carried out under the Egg Product Inspection Act (EPIA) as part of USDA’s Shell Egg Surveillance (SES)^67^. While successful in detecting these important food safety related pathogens, such surveillance completely overlooks the poultry health related pathogens-especially avian viral pathogens. We have demonstrated that egg (albumin) based screening and disease surveillance can be pretty effective in picking floating pathogens, and help us understand the disease burden, patterns and trends in general.

### Limitation of our study

This study was based on and relied upon the eggs being provided by the participating hatcheries. Most of these hatcheries only requested to have their eggs screened based on suspected clinical signs (and often after some mortality). Hence, they did not provide eggs routinely and regularly. Because of this, we were not able to establish the real disease occurrence trend across each hatchery nor were we able to tell whether the de-contamination efforts they made actually worked. Furthermore, randomly selecting 10% of the eggs might not have the sensitivity needed to pick all the pathogens in a given farm; we only screened a fraction of eggs in each batch due to cost consideration. We could have integrated an environmental screening and bio-security assessment as a part of a thorough disease surveillance system in poultry industry/farms, however, we were not able to do that in this study. Interestingly, we did observe some seasonal variability of disease occurrences-wet or rainy season harboring more pathogens than dry season. However, we need more data points to look into this further.

### Implications and Utility of Egg based Pathogen screening

Molecular detection of pathogens in egg albumin can provide important information to put together an early containment strategies for poultry farmers in particular and animal health efforts in general. It can be especially beneficial to hatcheries as they are often the contamination source-spreading disease from egg to day-old chicks, and eventually to the whole poultry production value chain. Egg-based disease screening can be an effective One Health surveillance tool as well, as it can pick up important Zoonotic pathogens such as Influenza A, and help stem pathogen spill-overs, thereby safeguarding human health. Albumin screening, as an early detection tool, can also assess biosecurity effectiveness in hatcheries and help curb horizontal and vertical transmission of avian diseases. In a developing country like Nepal, where resources are limited, easy to access pathogen screening samples like eggs and highly sensitive and accurate molecular (PCR) tools can help in building important avian disease surveillance tool. With the advent of next generation DNA sequencing and Genomics technology, we can even screen for a broader viral, bacterial and other pathogens using same (single) sample source. High throughput in data acquisition made possible by such new technologies certainly can make disease surveillance fast, easy and affordable. Our study is an initial step towards that direction.

## Acknowledgement

We would like to thank the participating hatcheries for their contribution in this important study. Our gratitude also goes to the experts from the Intrepid Nepal Pvt. Ltd. and the Center for Molecular Dynamics Nepal (CMDN) for their help with the Genomics analysis. And finally, a big thank you to the Poultry Association of Nepal for their continued support.

## Author Contributions

Conceptualization: Rajindra Napit

Data curation: Shreeya Sharma, Kavya Dhital, Sajani Ghaju, Prajwol Manandhar

Field support: Dhiraj Puri

Laboratory analysis: Kavya Dhital, Sajani Ghaju, Udaya Rajbhandari, Amit Basnet

Methodology: Rajindra Napit, Shreeya Sharma

GIS Map: Prajwol Manandhar

Supervision: Rajindra Napit, Dibesh Bikram Karmacharya

Validation: Dibesh Bikram Karmacharya

Writing – original draft: Shreeya Sharma

Writing – review & editing: Dibesh Bikram Karmacharya

